# Genome-wide Nucleosome Positioning and Associated Features uncovered with Interpretable Deep Residual Networks

**DOI:** 10.1101/2024.02.09.579668

**Authors:** Yosef Masoudi-Sobhanzadeh, Shuxiang Li, Yunhui Peng, Anna R Panchenko

**Affiliations:** Department of Pathology and Molecular Medicine, Queen’s University, Canada; Institute of Biophysics and Department of Physics, Central China Normal University, Wuhan, China; Department of Biology and Molecular Sciences, Queen’s University, Canada; School of Computing, Queen’s University, Canada; Ontario Institute of Cancer Research, Canada

**Keywords:** Interpretable deep learning, Nucleosome positioning, Nucleosome positioning features, ResNet

## Abstract

Nucleosomes represent elementary building units of eukaryotic chromosomes and consist of DNA wrapped around a histone octamer flanked by linker DNA segments. Nucleosomes are central in epigenetic pathways and their genomic positioning is associated with regulation of gene expression, DNA replication, DNA methylation and DNA repair, among other functions. Building on prior discoveries, that DNA sequences noticeably affect nucleosome positioning, our objective is to identify nucleosome positions and related features across entire genome. Here we introduce an interpretable framework based on the concepts of deep residual networks (NuPose). Trained on high-coverage human experimental MNase-seq data, NuPose is able to learn sequence and structural patterns and their dependencies associated with nucleosome organization in human genome. NuPoSe can be used to identify nucleosomal regions, not covered by experiments, and be applied to unseen data from different organisms and cell types. Our findings point to 43 informative DNA sequence features, most of them constitute tri-nucleotides, di-nucleotides and one tetra-nucleotide. Most features are significantly associated with the structural characteristics, namely, periodicity of nucleosomal DNA and its location with respect to a histone octamer. Importantly, we show that linker DNA features contribute ∼10% to the quality of the prediction model, which together with comprehensive training sets, deep-learning architecture and feature selection may explain the advanced performance of NuPose of 80-89% accuracy.

## Introduction

The human genome is packaged into chromatin with nucleosomes serving as building blocks. Nucleosomes consist of ∼147 base pairs (bp) of DNA (“nucleosomal DNA”) wrapping ∼1.7 turns around a histone octamer^1^. Nucleosomes are connected together by about 20-90 bp DNA segments, called linkers. There are 30 million nucleosomes in the human genome, and their specific positions are associated with key biological functions mediating DNA accessibility, gene expression, DNA methylation and binding of various chromatin factors^2–4^. Different experimental methods have been proposed for nucleosome mapping which include DNase-seq, ATAC-seq and NOME-seq, but they generally lack the single nucleotide precision which is necessary for deciphering sequence patterns modulating the preferable location of DNA on histone octamers^5,6^. Other techniques, employing directed chemical cleavage, may offer very high resolution but might not be viable for mapping across the entire genome^7^. MNase-seq is widely used method for nucleosome detection, despite its demand for extensive sequencing coverage^8,9^. For instance, to achieve a high-resolution map of nucleosomes in human cells, several billion reads are necessary. Despite the availability of extensive datasets, the experimental mapping efforts have not precisely determined nucleosome positions (NP) in the entire human genome which underscores the necessity to develop comprehensive prediction methods for NP.

Nucleosome positioning in cells in general is influenced by the DNA sequence features, by chromatin physical barriers and by chromatin factors which can slide nucleosomes^10–14^. Although DNA molecule is flexible, its certain conformations may be energetically favorable for nucleosome formation depending on its sequence^15–17^. Motivated by these ideas, various studies have been carried out to identify the DNA-specific features that are effective in forming nucleosomes^18–22^. Although first analyses on this subject were performed on very small data sets, these studies were capable to distinguish certain sequence patterns governing nucleosome positioning. It was proposed that the deformation cost of bending DNA around a histone octamer could depend on the locations of certain pyrimidine-purine dinucleotides that were easier deformed compared to others^23,24^. While there is no consensus on this topic, sequence patterns were described where A/T rich sequence motifs determined the rotational orientation of DNA so that minor grooves faced towards the histone octamer, whereas G/C rich motifs were associated with the minor grooves facing outward^25,26^. However, there is no consensus on the subject. Generally, the contribution of genomic sequence in explaining the *in vivo* nucleosome positioning has been proposed to be substantial^27,28^.

The experimental determination of NP is a time- and cost-consuming process. Hence, different discriminant and generative computational methods were proposed, inspired by the idea of NP dependence on DNA sequence^29–33^. One group of studies used di- and tri-nucleotide composition trying to maximize the distance in feature space between nucleosomal footprints and nucleosome free regions^34^. These approaches did not account for precise locations of DNA sequence patterns and this choice was partially explained by the relatively low-resolution and a small amount of data coming from the SELEX and microarray MNase-seq techniques. In the later studies, the authors tried to account for the DNA local structural properties and long-range sequence-order effects which were proven to be very effective^35^. Most recently, several deep learning (DL) approaches have been developed with the goal of predicting NP in a genome^36,37^. One such approach (LeNup) proposed to combine the concepts of the convolutional neural network (CNN), inception modules, and a gating mechanism^38^.

Despite all these above-mentioned efforts, the DNA sequence-based features effective in forming nucleosomes have not been well understood because of the following reasons. First, most machine learning studies have focused on predicting NP rather than identifying NP features. Second, existing NP prediction approaches have used only parts of the human genome for training due to the high memory and time complexities of these algorithms. For example, in CNN, numerous kernels in convolution and pooling layers do not allow to efficiently train the model and identify NP features in the on the whole genome scale^39^. Furthermore, given the large scale of NP data, using advanced deep learning techniques like capsule networks also poses a challenge in terms of time complexity^40,41^. Finally, existing studies have neglected to consider the linker DNA regions adjacent to nucleosomes.

Here, powered by the high coverage human data, we have developed an interpretable deep-learning framework based on the concepts of deep residual networks (ResNet) and a two-step feature selection approach. Our proposed method extracts a variety of NP features that are not typically extractable by common deep-learning approaches. These features undergo a two-step feature selection process, and the final NP prediction model and features are selected using a ResNet-based approach. We find 43 sequence features that are important for identification of nucleosome regions genome-wide with the accuracy up to 89%. Importantly, we show that not only nucleosomal but also flanking linker DNA sequence is important for nuclesome positioning and in total contribute ∼10% to the quality of the model.

## Materials and Methods

Figure 1 shows the main steps of our computational framework (NuPose) which can score the nucleosome positioning (dyad locations) genome-wide and evaluates the importance of features contributing to the quality of prediction model. NuPose includes the following major steps: (a) identification of nucleosome dyad from the experimental MNase-seq data, (b) feature extraction, (c) generation of candidate subsets of features, and (d) selection of the most effective features in forming a nucleosome and generating an NP prediction model.

**Figure 1.**
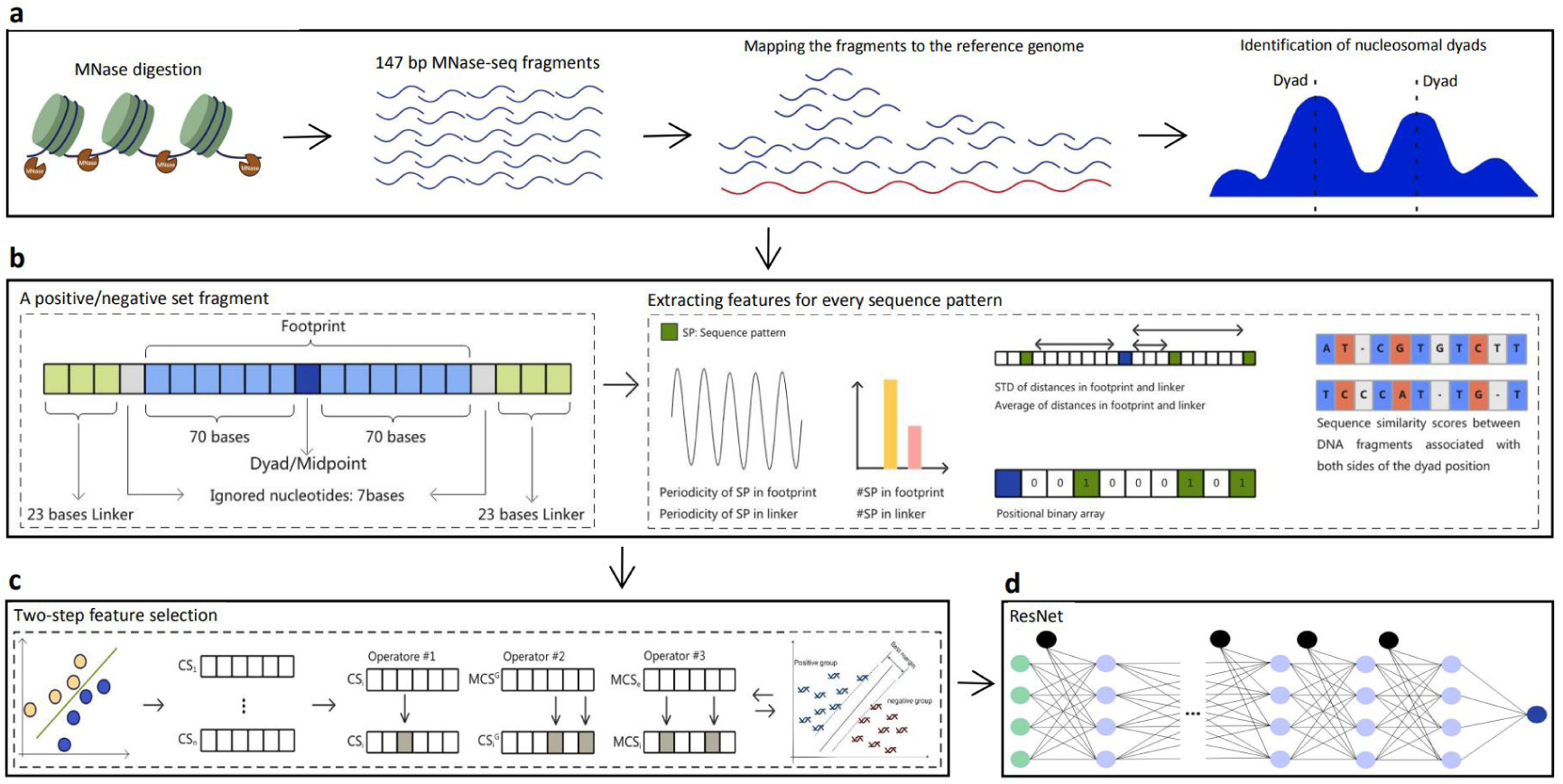
The framework of the proposed deep learning method (NuPose). (a) Mapping of MNase-seq fragments to the reference genome and identification of nucleosomal dyads. (b) Extraction of five groups of features from nucleosomal and inter-nucleosomal regions. (c) Feature selection approach to generate candidate feature subsets. (d) Selection of nucleosome positioning features and generation of prediction model using ResNet.

### Identifying nucleosome positions from the MNase-seq data

We followed the protocol developed in our previous study^42^. High-coverage data of paired-end 147 bp length MNase-seq fragments from seven human lymphoblastoid cell lines (GSE36979) were used in this study for mapping of nucleosome positions^28^. Fragments of 147 bp lengths are the most abundant in this data set and correspond to the nucleosomal DNA, whereas longer or shorter fragments may result from over- or under-digestion of nucleosomal DNA, can come from the sequence-specific spontaneous DNA unwrapping and may provide less precise estimates of individual nucleosome positions. The fragments (both complementary strands were used) were mapped to the reference human genome (GRCh37). Since there can be multiple overlapping nucleosomal fragments, we applied a previously developed protocol to identify the representative dyad positions^43^. First, the fragments’ midpoint counts (dyad counts) at each genomic location were smoothed out (Eq. 1):

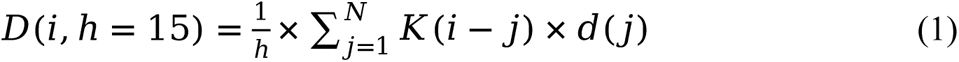

where *N* is the length of a chromosome, and *d(j)* indicates the total number of mapped DNA fragments with their midpoint nucleotides placed at the j^th^ genomic location. Here, *K* is the tri-weight kernel function (Eq. 2), and *h* is the bandwidth of the kernel function. The value of *h* was tuned up to 15, as prior studies have shown that smaller values for *h* can increase the accuracy of identifying NP^28,43^.

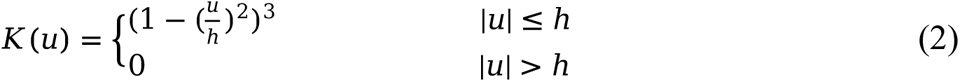

Second, the bwtool software package was used to obtain the local maximum values of the smoothed dyad counts^44^. The minimum distance between the neighboring local maxima was set to 150 bp with “find local-extrema -maxima -min- sep=150”. Then, for every 60-bp window centered at each local maxima, the dyad location with the highest number of dyad counts was determined as the representative dyad and any other dyad positions were discarded. In the cases where two or more dyad positions had the same dyad counts within the same interval, the representative dyad was determined as the one located closest to the local maximum of the smoothed counts.

As a result, we were able to determine nucleosome positions for over 1 million nucleosomes, from which the positive and negative data sets were chosen. For a positive set, 100 bp long fragments were extracted from both sides of the dyad position (Figure 1b), with 73 bp on each side of the dyad representing the *nucleosomal footprint*, and the remaining 27 bp on each side representing the *linker* DNA. The negative data set was taken from the *inter-nucleosomal* regions that did not overlap with any nucleosomal DNA fragments. For a negative set, we identified the midpoint between two consequent nucleosomal dyads, separated by >400 bp distance, and then included 100 bp on both sides from the midpoint. Both positive and negative datasets included genomic regions of 201 bp lengths (so called *samples*). In generating datasets, to avoid biasing towards sequence repeats, the repeated genomic regions were disregarded using the WindowMasker software tool^45^. As a result, each of the positive and negative datasets comprised 115,640 one-stranded DNA sequence regions (a total of 231,280).

### Extracting features

For every DNA strand from positive and negative data sets, a total of 4^2^+4^3^+4^4^=336 di-nucleotide, tri-nucleotide and tetra-nucleotide sequence patterns were defined, and for every sequence pattern, five groups of features were extracted (Figure 1b). The first group pertained to the total number of occurrences of a pattern within a sequence. The second group corresponded to statistical features, the average and standard deviation values of the distribution of distances between a pattern’s occurrences and the nucleosomal dyad. For the negative set, distances were measured from the midpoint of the inter-nucleosomal region (counted in the number of bases). Previous studies demonstrated that the frequency of di-nucleotides across nucleosomal DNA follows 10-11 base pair and, in some cases, 12 base pair periodic patterns. Therefore, we introduced the third group of features, calculated as the total number of occurrences of a sequence pattern separated by multiples of k (k = 10, 11, or 12 bases):

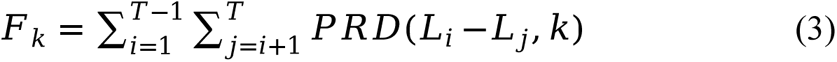

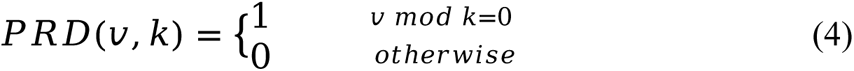

where *L_i_* is the genomic location of a sequence pattern and *T* is the total number of occurrences of a sequence pattern in a data sample. Due to the two-fold symmetry of DNA in nucleosome structures, DNA complementary strands should be structurally superimposed if the nucleosome structure is rotated by 180 degrees. Considering the symmetry attribute of a nucleosome, the fourth group of features included sequence similarity scores calculated. Finally, the fifth group pertained to the presence/absence of a certain sequence pattern at a certain distance from the dyad position (a midpoint in case of a negative set). All features were extracted for each of the 336 sequence patterns in every data sample. Moreover, since micrococcal nuclease has a strong sequence specificity and cleaves the 5’ side of A/T much faster than of G/C at flanking regions, we ignored regions located at 71-77 bases away from the dyad. All in all, a total of 34,276 features were defined and extracted for every data sample.

### Selecting informative features

Selecting an optimal subset of features is computationally intensive (a non-deterministic polynomial-hard problem), so it is not practical to score all possible subsets of features^46^. We introduced an interpretable multi-step feature selection approach and generated an NP prediction model based on the concepts of ResNet. In the first feature selection step, the Pearson correlation coefficients (PCC) were calculated between all feature values, and as a result 47 features were found to be redundant with PCC greater than 0.5 when compared to other features. The remaining features underwent the second step of the feature selection, which involved combining our previously developed metaheuristic *Trader* algorithm and support vector machine (SVM) classifier (we refer to it as SVM^Trader^)^47^. *Trader* speeds up the search by updating each feature subset by introducing random changes to candidate solutions (CS) to explore new regions of the global search space and aiding in escaping from the local minima. Additionally, the algorithm divides candidate solutions into several groups, enabling diversification of the search. This process allows for independent and more effective exploration of different regions within the global search space. Consequently, it leads to faster convergence and improved performance as was shown in the previous study^47,48^. The pseudo code of the *Trader*-based feature selection method is presented in the supplementary file.

At the end of the above-described feature selection step, a total of 100 improved CSs were generated and analyzed using the fully connected part of NuPose, which was designed based on the concepts of ResNet^49^. NuPose’s architecture included 50 fully connected hidden layers and differed from the architecture of ResNet by replacing convolution and pooling layers with the feature extraction and selection procedure described above (Figure 1 and Figure S1). Skip connections feed the input of a residual block to the output of another residual block (Eq. 5), with each block consisting of five fully connected hidden layers. These skip connections help to minimize the vanishing gradient problem and increase the performance of the deep neural network.

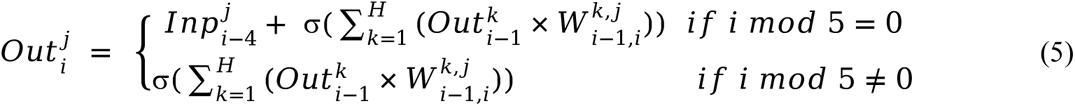

where 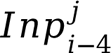 denotes the input of the *j^th^* neuron associated with the *i-4^th^* layer, and *H* is the total number of neurons in a hidden layer. 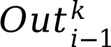 refers to the output of the *k^th^* neuron associated with the *(i-1)^th^* layer, and 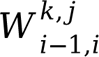 is the weight of the edge connecting the *k^th^* neuron of the *(i-1)^th^* layer to the *j^th^* neuron of the *i^th^* layer, respectively. Here, σ is an activation function of a layer *i*, in our case the sigmoid function. The epoch number and batch size parameters were set to 10,000 and 500, respectively. To prevent overfitting, the early stopping technique was used with a value of 5 for the patience parameter. The ResNet described above was applied to each of the 100 improved CS and certain NP features were selected based on the weights associated with the first layer of the described ResNet, similar to a previously introduced DL method^50^. Features with the weights larger than the mean value of all weights (*µ*) associated with edges connecting neurons from the first layer to those of the second layer were chosen

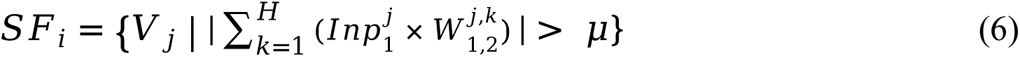

where *SF_i_* and *V_j_* represent a set of selected features from the *i*^th^ CS and *j*^th^ features within that CS, respectively. Here, *Inp^j^* is associated with the value of the *j*^th^ feature.

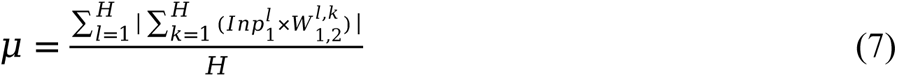

The union of the selected features from each CS was chosen as *informative features* in the formation of a nucleosome 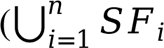, where *n* is the total number of CSs), and the final NP prediction model was generated based on these informative features. The informative NP features were then ranked based on their influence on the NuPose model. For this purpose, several NP prediction models were generated by removing each of the informative features and keeping the remaining ones. The areas under the receiver operating characteristic curves (AUC-ROC) were then compared using the DeLong test^51^. If Z-score was greater than 1.96, then the two NP models were assumed to significantly differ from each other with p-value <0.05^52^. The final ranking of features was determined by the inverse order of the DeLong test Z-scores since a significant drop in the Z-score of the model performance upon removing a feature would correspond to the most informative feature in the final ranking. The NuPose workflow was implemented in the Python programming language equipped with the Keras deep learning library.

## Results

### Comparing NuPose with previous methods

In this section, we compare NuPose performance with other previously introduced NP prediction methods^38,53–55^. These methods have been developed and evaluated based on the main three datasets: (i) derived from *H.sapiens* containing 2,273 positive and 2,300 negative samples (data set *HS*), (ii) derived from *C.elegans* consisting of 2,567 positive and 2,608 negative samples (data set *CE*), and (iii) *D.melanogaster* comprising 2,900 positive and 2,850 negative samples (data set *DM*)^56^. It should be mentioned that due to the computational complexity, all these studies trained their prediction models on a relatively small fraction of genomes (50 times smaller than a data set used to train NuPose), for example, only chromosome 20 was used in case of *H.sapiens (HS)* dataset. Besides, these datasets consisted of 147-base DNA sequences and excluded linker DNA regions. Therefore, we re-trained NuPose on these datasets without considering linker regions (so called NuPose*). Consequently, to directly compare NuPose* performance with the results reported in other methods’ papers, we have generated predictions using 5-fold and 20-fold cross-validation (Table 1). As can be seen in this table, the NuPose* and LeNup methods outperformed other techniques in terms of various performance criteria. Moreover, even though NuPose* model did not used the linker features, it surpassed LeNup by ∼8% and 2% in terms of MCC for the *H.sapiens (HS)* and *C.elegans (CE)* datasets respectively, whereas for *D.melanogaster (DE)* dataset LeNup outperformed NuPose by 2%.

**Table 1.**
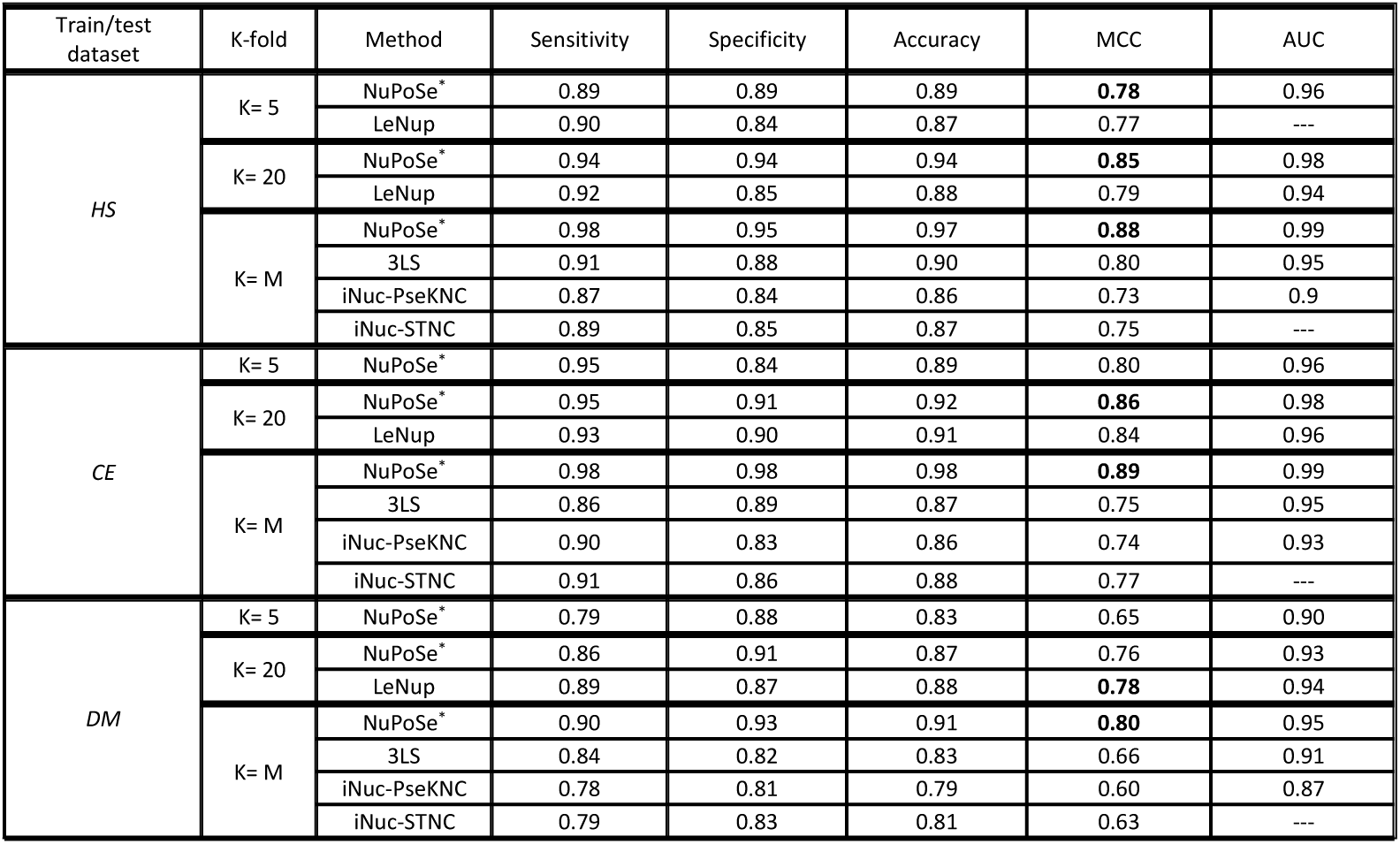
Comparing the performance of NuPoSe* with previous studies. Values in bold show the best-acquired result for a given organism and cross-validation method. *M* indicates the total number of sequences in the dataset. NuPoSe*corresponds to NuPoSe trained on datasets without linker DNA, since previous studies did not include linkers into consideration. HS, CE and DM refer to data from *H.sapiens*, *C.elegans* and *D.melanogaster*, respectively.

### Generalization of the model across different species and cell lines

Next, we asked, if NuPose framework trained on one specific cell line can be applicable for identifying NP in a different cell line in the same organism. To answer this question, an additional MNase-seq dataset from CD4+T cells was obtained^56^. This dataset covered regions of the human genome in activated and resting CD4+T cells. We trained three models on our original human lymphoblastoid cell lines: SVM^Trader^, NuPose and a version of NuPose which excluded skip connections (NuPose^WS^, to evaluate the effects of skip connections on the performance of the NP prediction model). Then we applied these models to predict nucleosome positioning of CD4+T cells (Figure 2a). The results indicated a decrease in the performance of all models, especially SVM^Trader^, when applied to an independent set of CD4+T cells (Figure S2, S3), which is likely due to the fact that prediction models were trained on a set from a different cell line. However, NuPose performed quite well even though it was applied to a different cell type (AUC dropped from 0.95 to 0.87). Moreover, NuPose outperformed two other prediction methods on both resting and activated cells in terms of criteria shown in (Figure 2b and Figure S2). Interestingly, predictions produced by all three models were considerably more accurate for resting cells, compared to activated cells. Possible reasons explaining this result will be discussed later.

**Figure 2.**
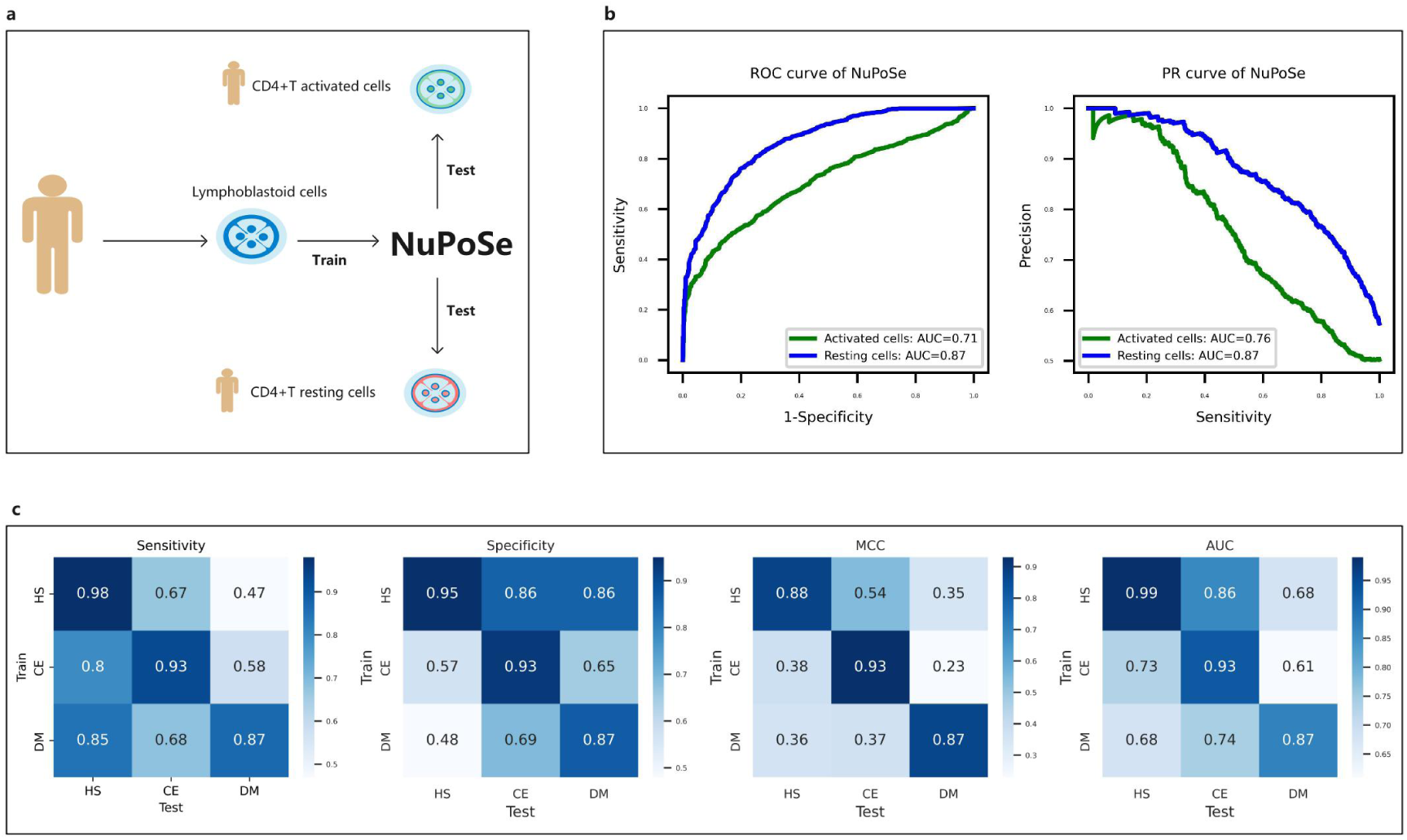
Generalization of the NuPose prediction model across different cell lines and organisms. (a-b) An overview and the performance accuracy of NuPose trained on the MNase-seq data from human lymphoblastoid cell lines and tested on the independent MNase-seq data derived from CD4+T cells: activating and resting cells. (c) Outcomes of models trained using data from one organism and applied to other organisms. HS, CE and DM refer to the data sets from *H.sapiens*, *C.elegans* and *D.melanogaster*, respectively.

Furthermore, we have assessed the ability of the model trained on one organism to predict nucleosome positioning in another organism. The results showed that NuPose* trained on the *HS* dataset outperformed models trained on *DM* and *CE* in terms of criteria shown in Figure 2c. Even though the performance dropped upon switching to a different organism, model generated based on the *HS* data reasonably predicted NP in *CE.* However, the prediction quality for *DM* using methods trained on other organisms was relatively low.

### Identification of informative nucleosome positioning features

First, we checked whether the inclusion of linker DNA features enhanced the prediction accuracy of our model. Therefore, we have developed a NuPose approach using two different schemes with (NuPose) and without the linker DNA-related features (NuPose*) (Figure 3a). Our results demonstrated that the linker DNA features resulted in more than 10% increase in the performance (MCC = 0.79 compared to 0.60 and AUC = 0.95 compared to 0.87), no matter what data representation, method or network architecture was used (Figure 3b, Table S1). It was observed that NuPose surpassed all other approaches, emphasizing the role of DL, skip connections and feature selection in increasing the performance (Figure S3 and Table S1).

**Figure 3.**
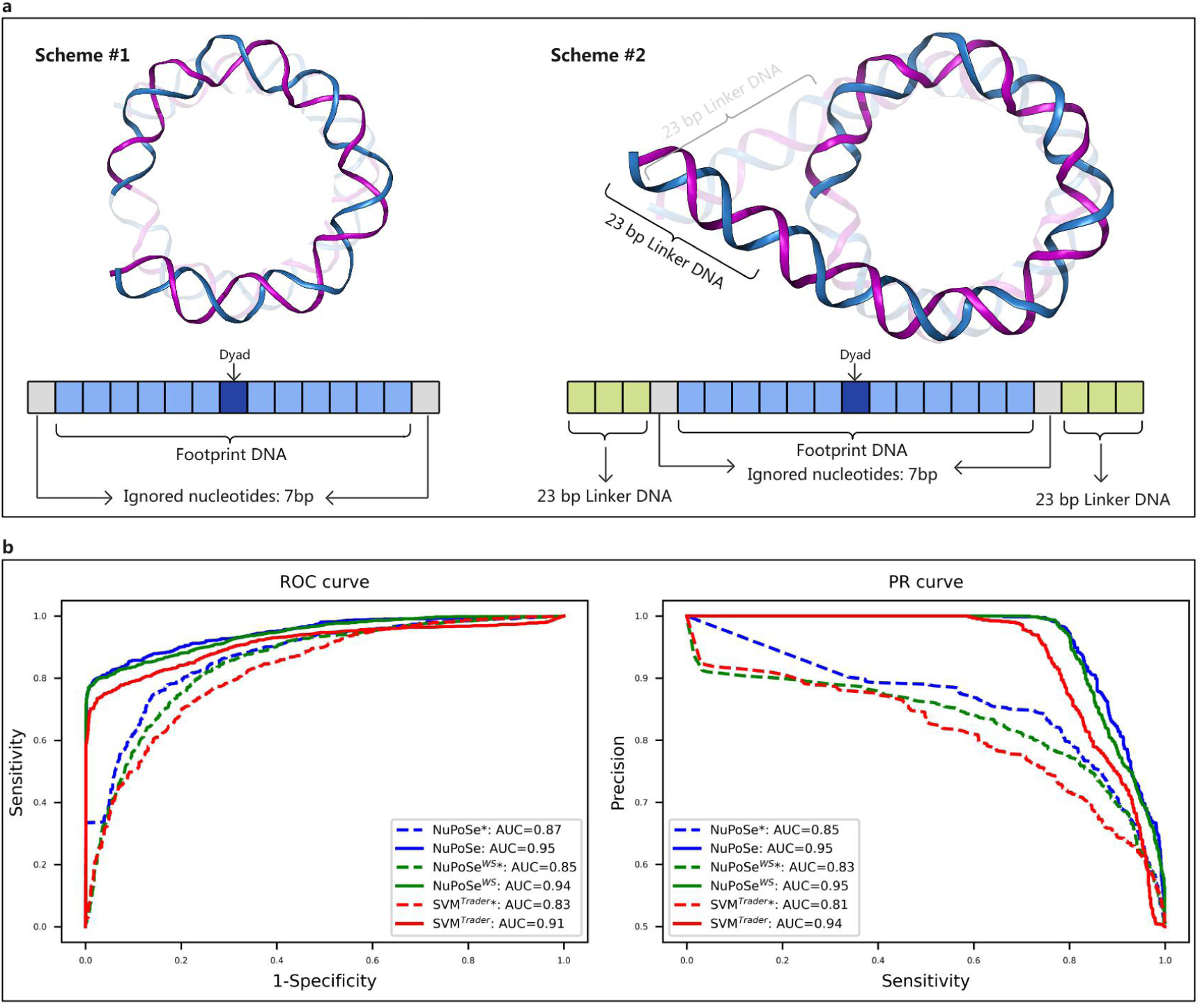
Enhancement of prediction accuracy by inclusion of linker DNA associated features. Two schemes were employed to generate datasets. (a) Scheme #1 comprises features derived from nucleosomal footprint DNA only, while scheme #2 considers both the footprint and linker DNA. (b) The ROC and PR curves of the methods. Dashed lines and asterisks (*) correspond to models which do not use linker DNA-related features. NuPoseWS is a version of NuPose which excluded skip connections. SVMTrader is a combination of Trader and SVM.

Next, we inquired about sequence features which had the most significant impact on the model’s performance. To assess the significance of each feature’s contribution to the quality of the model, we followed the protocol described in Methods. We showed that the second and fourth categories of features, (statistical and sequence similarity-based features) were not selected as being informative. Our results pointed to 43 informative features which governed the NP genome-wide (Figure S4, Tables S2-S5). We have also calculated the correlation between the informative features chosen by NuPose and those features that were not chosen by NuPose (Figure S5, S6). Among them, nine features were related to the periodicity of sequence patterns in the nucleosomal footprint DNA (Figure 4a and Figure S7, Table S3). Although we formulated our feature periodicity calculations in real space, the power spectra from the Fast Fourier Transform (FFT) confirmed the largest amplitude for the components with 10-12 bp periodicity (Figure S8). Based on the feature ranking results, the periodicity related feature, as a group was more influential than other types of features in distinguishing nucleosomal and inter-nucleosomal DNAs, contributing up to ∼40% to the quality of the model (Table S2 and Figure S4).

**Figure 4.**
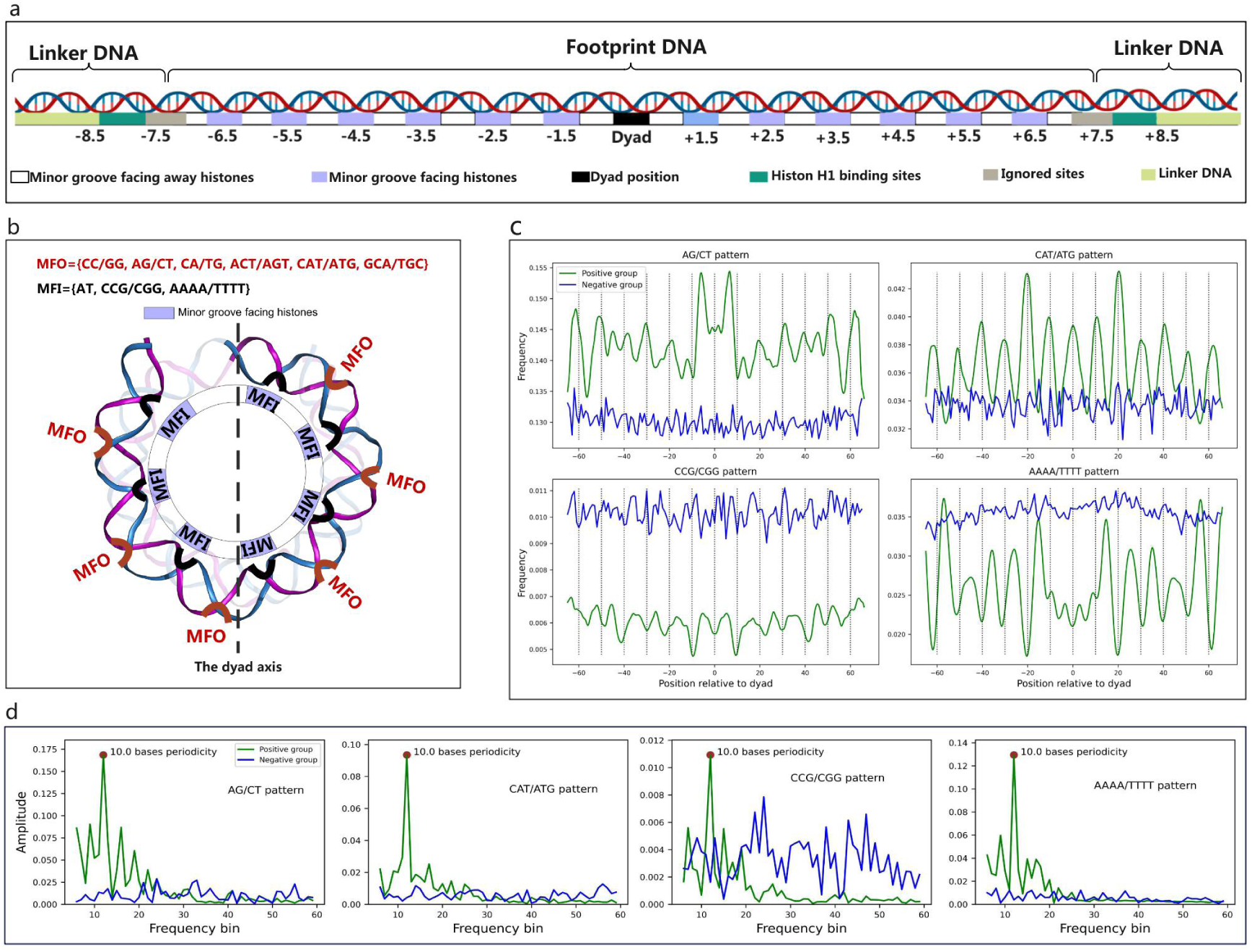
Illustration of the informative periodicity-related features. (a) Locations of DNA minor grooves facing a histone octamer (blue, MFI) and facing out (white, MFO) on nucleosomal DNA sequence. Integer numbers along the DNA sequence show the assignment of super-helical locations (SHL). (b) Illustration of the informative periodicity related features with statistically significant associations with MFO and MFI throughout the whole nucleosomal footprint (Table S3). Features associated with the specific SHL locations are shown in Figure 5. Highly correlated features with PCC >0.5 are AT and TA, CC/GG and CCA/TGG, CCG/GCC and GC but only one informative feature from the group is shown. (c) The frequency (#sequence patterns/#samples) of several representative periodicity-related informative features. Figures S5 and S6 show the results for all features. The dashed lines show minor grooves facing outward. (d) Results of the Fast Fourier Transform analyses. A component with the largest amplitude is shown as a red dot.

Out of all informative features, the most informative one was related to the CCG/CGG sequence pattern. Although this pattern was depleted in terms of its occurrences on nucleosomal compared to inter-nucleosomal regions, nucleosomal DNA had higher percentage of these sequence patterns separated by ∼10 base pairs (75% versus 55%). Other periodicity related features with high contribution to the quality of the model included tri-nucleotides AGT/ACT and CAT/ATG (∼11% in total) and dinucleotides AG/CT and AT (as well as TA due to its high PCC value with AT, Figure S5). The only informative tetra-nucleotide was AAAA/TTTT. Overall, our results indicated that the sequence patterns, whose periodicity has been selected as informative features, generally exhibited a much higher nucleosome occupancy values relative to other sequence patterns (Figure S9).

Powered by the structural knowledge of the locations of nucleosomal DNA with respect to the histone octamer, we calculated the enrichment and depletion of different sequence patterns with respect to the DNA minor groove facing histone octamer (MFI) (corresponding major grove will be facing out) or a minor groove facing outward (MFO) and corresponding major groove will facing a histone octamer. (Table S3). We found that AT, CCG/CGG and AAAA/TTTT patterns were enriched on minor grooves facing inward with high effect size (Odds ratio = 2.17 - 4.38). It is consistent with previous experimental measurements of tri-nucleotide bendability showing that these patterns have low bendability toward the major groove (and high bendability toward minor grooves)^57^. Interestingly, periodicity of AAAA/TTTT (an example of an A-tract) is an informative factor in NP prediction, even though A-tracts are more rare in nucleosomal compared to inter-nucleosomal DNA, and long A-tracts (longer than 5 bp) may serve as excluding nucleosome boundaries^58,59^. On the other hand, all other periodicity related features were enriched on minor grooves facing out (Table S3).

Among other informative features, 12 features were associated with the total number of occurrences (composition) of certain di- and tri-nucleotide patterns (Figure 5a). Nucleosomal footprints regions were overall depleted in terms of A/T-based di- and tri-nucleotide sequence patterns compared to the inter-nucleosomal regions, whereas linker regions were enriched with respect to AT and TA dinucleotides compared to the inter-nucleosomal regions (Table S4, Figure S6).

**Figure 5.**
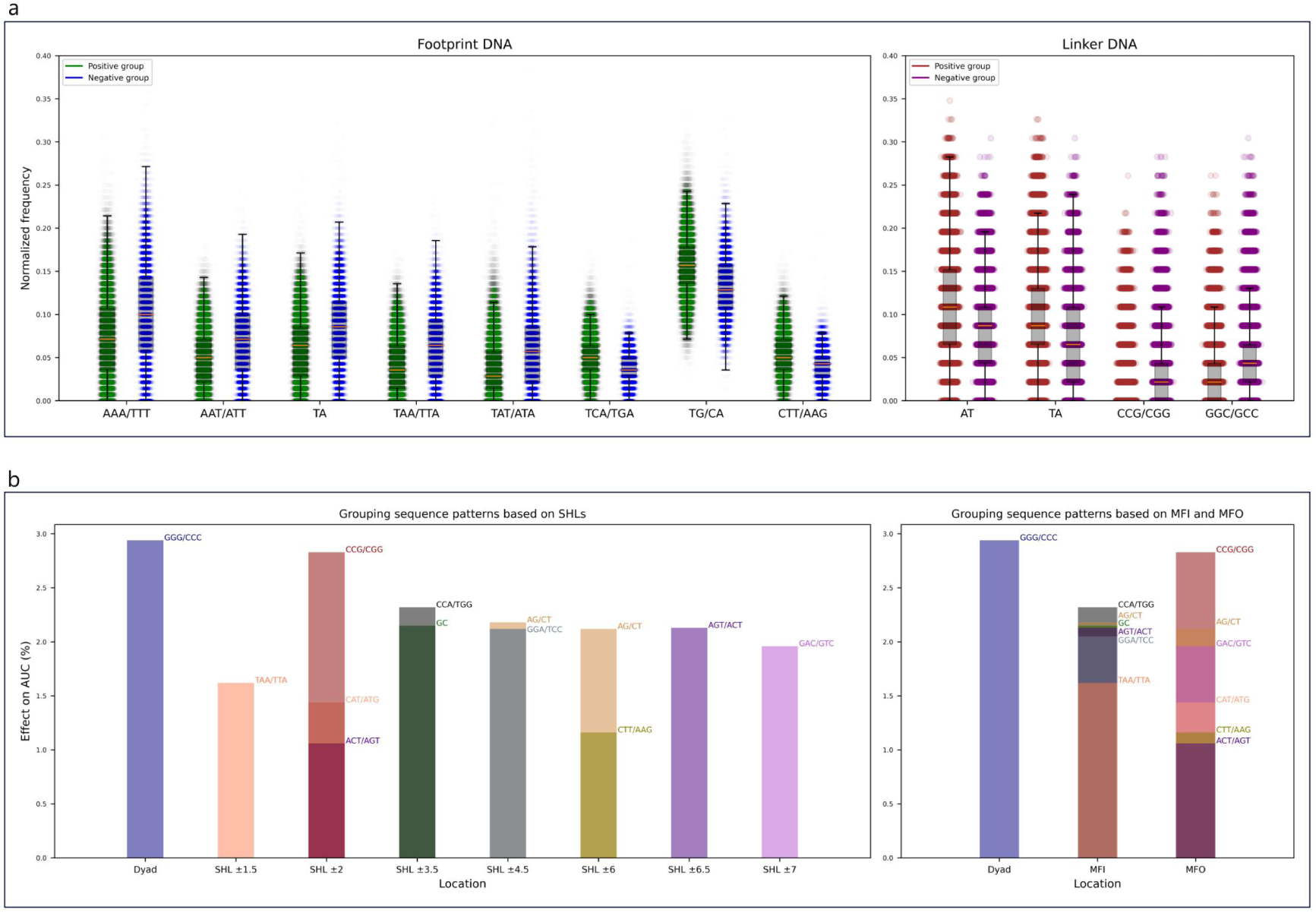
The composition and distance-related informative features. (a) Features associated with the composition of di- and tri-nucleotide sequence patterns in positive (green and brown) and negative (blue and purple) groups. Assignments to super-helical locations (SHL) for nucleosomal footprints were based on Figure 4a. Y-axis shows the frequency of a given sequence pattern from Figure 4c divided by the length of the nucleosomal footprint (left panel) or linker DNA (right panel) for comparison. (b) The informative distance-related features were grouped based on SHL (left) and MFI/MFO (right). Y-axis shows the percentage of AUC drop when a specific feature is removed. Note that the features may be depleted or enriched at a given location (Table S5).

Finally, the remaining 22 features were associated with the presence of specific sequence patterns at certain positions relative to the dyad (Figure S10 and Table S5). Each of these features contributed only up to 3% to the quality of the model individually and majority of them had statistically significant positive or neagtive associations with the structural characteristics of nucleosomes. For example, highly informative CCG/CGG, ACT/AGT, and CAT/ATG, were associated with SHL ±2 (Figure 5b), where the CCG/CGG sequence pattern was depleted at this location (minor groove facing out, MFO), while the latter two were enriched. Two sequence patterns AG/CT (and AGT/ACT) were enriched at position SHL ±6 (facing out) and depleted at SHL ±4.5 (facing in), consistent with its propensity to be enriched on the minor grooves facing out. Interestingly, some linker-associated features were also position specific (Table S5).

## Discussion

Powered by the idea of dependency of nucleosome positioning on DNA sequence, we have introduced an interpretable deep residual network-based framework for selecting informative NP features and classifying nucleosome regions across human genome. Our method showed superior performance compared to previous approaches that could be due to the use of deep-learning architecture, extensive training set and feature selection algorithms. We have confirmed that our method can distinguish nucleosomal from inter-nucleosomal regions in organisms other than human, although the performance may drop depending on the organism and cell type. In addition, our analysis allowed us to gain insights into the contribution of *in vivo* factors into nucleosome positioning. Namely, we showed that in activated cells, where *in vivo* factors may actively modify chromatin structure and affect nucleosome positioning, the accuracy of our model decreased to ∼60% compared to ∼80% in resting cells. This is in line with the previous study that demonstrated that a pronounced similarity between the *in vivo* and *in vitro* nucleosome maps diminishes in the regions of actively expressed genes^5^.

A total of 43 features were identified as informative and ranked based on their relevance. Many previous studies identified di-nucleotide periodic preferences (WW/SS/YY/RR) that might influence nucleosome positioning, where W, S, Y, and R respectively represent A/T, C/G, C/T, and A/G^16,60^. We have confirmed this trend for AT, TA, CA/TG, CC/GG and AG/CT dinucleotides, but showed that the vast majority of informative features constituted tri-nucleotides, some of them were extensions of the informative di-nucleotide features: (C)AT/AT(G), (G)CA/TG(C), CC(G)/(C)GG, AG(T)/(A)CT, and AAA(A)/(T)TTT (a nucleotide in parentheses indicates the extension of the di-nucleotide). Indeed, di-nucleotide based models do not fully capture the nucleosome formation as bendability of di-nucleotides depends on the local sequence context, and physico-chemical properties of longer sequence patterns are not additive of corresponding base pair steps^57,61^. Prior studies also argued that the 10-11 bp periodicity of the DNA sequence patterns observed in nucleosomes was related to the DNA bendability in those regions facing a histone octamer^62,63^. Therefore, we have estimated the statistical power of the association of informative features with the DNA and nucleosome structural attributes. Indeed, the majority of informative features demonstrated statistically significant associations with the minor and major DNA grooves and their locations with respect to a histone octamer. The most informative CCG/CGG feature tends to be located at a major groove facing outward (minor groove inward). We hypothesize that it could be associated with the 5’ cytosine methylation which always happens in the major groove, which, in turn, should be solvent exposed. Indeed, DNMT1 methyltransferases have strong specificity for CCG patterns and there are multiple studies showing that DNA methylation is favoring nucleosome formation, although in the genomic context dependent fashion^64–67^.

It should be mentioned that our model is based not only on the preference of certain sequence patterns, but also on their depletion. For example, we have confirmed that nucleosomal footprints were depleted in terms of A/T containing nucleotides compared to inter-nucleosomal regions, whereas nucleosome flanking linker regions were enriched with A/T-based sequence patterns. It can be attributed to the requirement of the DNA flexibility of linker regions, their potential binding to H1 histone, as well as the formation of the stem-like structures^68^. In line with this, we showed that the linker DNA features lead to ∼10% increase in the performance. This is important as prior nucleosome positioning prediction methods completely overlooked the effect of linkers. Indeed, nucleosomes can occupy different alternative positions but not too far from their original ones, namely, in the vicinity of the linker regions. In fact, it was shown previously that periodic oscillation of sequence patterns may go beyond the ends of nucleosomes^68^.

Finally, we showed that the distance of sequence patterns from the dyad can also be important. When a histone octamer binds to DNA, it confronts a choice among various potential translational positions determined by the mechanical properties of DNA, its shape parameters and by the affinity of histone-DNA interactions. The interactions between histones and DNA within nucleosomes exhibit diverse strengths and may play distinct roles in bending the DNA and influencing the nucleosome stability^69,70^. In addition, one could posit that features dependent on location may arise due to the need for sequence-specific recognition of nucleosomes by various binding partners, rather than being driven solely by the necessity for DNA bending. For example, it was shown previously that pioneer transcription factors and other nucleosome binders use distinct binding modes recognizing specific regions of nucleosomal DNA^42,71–73^.

Our results do not imply that the identified features are the sole influential factors in forming nucleosomes. It is imperative to emphasize that sequence-governed nucleosome positioning is statistical in nature, and the informative features do not contribute to nucleosome positioning individually, as they might not appear in all genomic regions at the same time. In other words, diverse combinations of the identified features (2^43^ possible states) may correspond to different nucleosomes with certain genomic location preferences, as shown earlier^74–76^. The application of our model is not confined only to understanding the mechanisms of nucleosome positioning. It has been shown that nucleosome positioning can be used as a feature in classifying disease subtypes in cancer and in improving the sensitivity of liquid biopsy^77^.

## Abbreviations

ACC: Accuracy
AUC: Area under curve
CNN: Convolutional neural network
CS: Candidate solution
DL: Deep learning
FFT: Fast Fourier Transform
NP: Nucleosome positioning
MCC: Matthew’s correlation coefficient
MFI: Minor groove facing histone octamer
MFO: Minor groove facing outward from the histone octamer
NuPose: prediction model/method trained on all features
NuPose*: NuPose trained without flanking DNA linkers
NuPose^WS^: A version of NuPose with excluded skip connections
PCC: Pearson correlation coefficient
PRE: Precision
ResNet: Deep residual network
ROC: Receiver operating characteristic SEN: Sensitivity
SHL: Super-helical location SPC: specificity
SVM: Support vector machine
SVM^Trader^: A combination of *Trader* and SVM

## Acknowledgments

The authors thank Victor Zhurkin for comments on the manuscript. Y.M.S., S.L. and A.R.P. were supported by the Department of Pathology and Molecular Medicine, Queen’s University, Canada. Y.P. was supported by the National Natural Science Foundation of China (No.12205112) and Fundamental Research Funds for Central China Normal University. A.R.P. acknowledges the support of the New Frontier in Research Fund (NFRF) and the Natural Sciences and Engineering Research Council of Canada (NSERC). A.R.P. is the recipient of a Senior Canada Research Chair in Computational Biology and Biophysics. This study was conducted with the support of the Ontario Institute for Cancer Research through funding provided by the Government of Ontario. The views expressed in the publication are the views of the authors and do not necessarily reflect those of the Government of Ontario.

## Code availability

The implemented source codes have been deployed in a GitHub repository accessible through the following link: https://github.com/Panchenko-Lab/NuPoSe

